# Phylogeography and Proline amino acid usage of Asian tiger mosquito *Aedes albopictus* (Skuse 1894) populations along landscape gradients in Indonesia

**DOI:** 10.1101/2021.03.14.435316

**Authors:** Andri Wibowo

## Abstract

Asian tiger mosquito *Aedes albopictus* (Skuse 1894) is one of mosquito-borne vector that globally distributed including Southeast Asia. Despite its wide distributions, information on phylogeography and its determinant factors influencing *A. albopictus* genetic diversity and speciation is still lacking, mainly in SE Asia country. Considering this, the phylogeography of *A. albopictus* along with its genetic attributes in Indonesia has been assessed. The results identify that there are 2 distinct clades of *A. albopictus* phylogeography. First clade was related to the human-mediated dissemination. Relative Synonymous Codon Usage (RSCU) shows the TTG, CTG, GTG, ACG, GCG, and GGG codons were seldom represented. While GGA, CGA, and followed by TAT, CAT, CCT, TCT, AAC, and TTC codons were common. From a total of 60 codons, 40% codon has RSCU > 1and 21.6 % has RSCU < 1. The differences between west and east part populations can be observed in proline amino acid signaled by CCT and CCC codons. In this amino acid, east *A. albopictus* has higher CCC than CCT codons. Since proline is functioned to provide energy for flight, then the differences of this proline related codon among *A. albopictus* populations were related to the landscape variations in west and east parts of Indonesia with east parts have more rugged landscape and this condition is quite demanding for aerial animal distribution since it requires more flight energy.

## INTRODUCTION

Mosquito-borne arboviral diseases have become a major global health concern. One of the mosquito-borne disease from *Aedes,* Dengue is the most important *Aedes*-borne viral disease in the world, causing 10,000-20,000 deaths per year) and almost 400 million new infections are estimated to occur annually. Arboviruses like Dengue, Zika and Yellow fever have spread worldwide following the expansion of their main vector *Aedes.* This mosquito has originated in Africa from an ancestral sylvatic and more zoophilic form *Aedes,* which expanded from tropical forests to urban areas giving rise to a domestic and anthropophilic form known as *Aedes*. This form was the only that succeeded in invading the rest of the world, forming a monophyletic group. *Aedes* arrived to the New World together with the first Europeans and Africans during the historical trans-Atlantic shipping traffic between 1500s-1700s, followed by the first reports of yellow fever and Dengue in the region (Powell & Tabachnick 2013).

One of important *Aedes* species is *A. albopictus* (Skuse 1894). This species that is also known as Asian tiger mosquito has been described as one of the 100 worst invasive species in the world. *A. albopictus* is originated from South and East Asia and this species has spread throughout the world mostly since the second half of the twentieth century, and it is now found on every continent except Antarctica (Kraemer et al. 2015). The new areas colonized by *A. albopictus* are ranging from disparate environments in tropical South America, Africa to the mostly temperate areas of Northern America and Europe. Tropical countries in Asia such as Southeast Asia are included in the endemic areas of Dengue and *A. albopictus,* one of which is Indonesia (Andriani 2016, Heriawati et al. 2020). WHO noted Indonesia as a country with the highest dengue cases in Southeast Asia and has high abundance of *A. albopictus*. According to Ministry of Health (Kemenkes 2016) in 2014-2016, there were 201,885 cases of Dengue fever spread throughout the province, with 77.96 Incidence Rate (IR) or morbidity rate per 100,000 populations, 0.79% of the case fatality or death rate and there were 1,585 cases caused death.. Regarding the magnitude of *A. albopictus* cases, whereas the information on genetic diversity of *A. albopictus* is still limited. While the genetic information of this vector is very important since it is needed for achieving effective */*.league cases eradications. This condition is the central issue of this study.

## MATERIALS AND METHODS

### Study area

This study was conducted in 3 locations across Indonesia region that are divided proportionally. Those locations included west, central and east parts of Indonesia (Figure 1).

**Figure 1.**
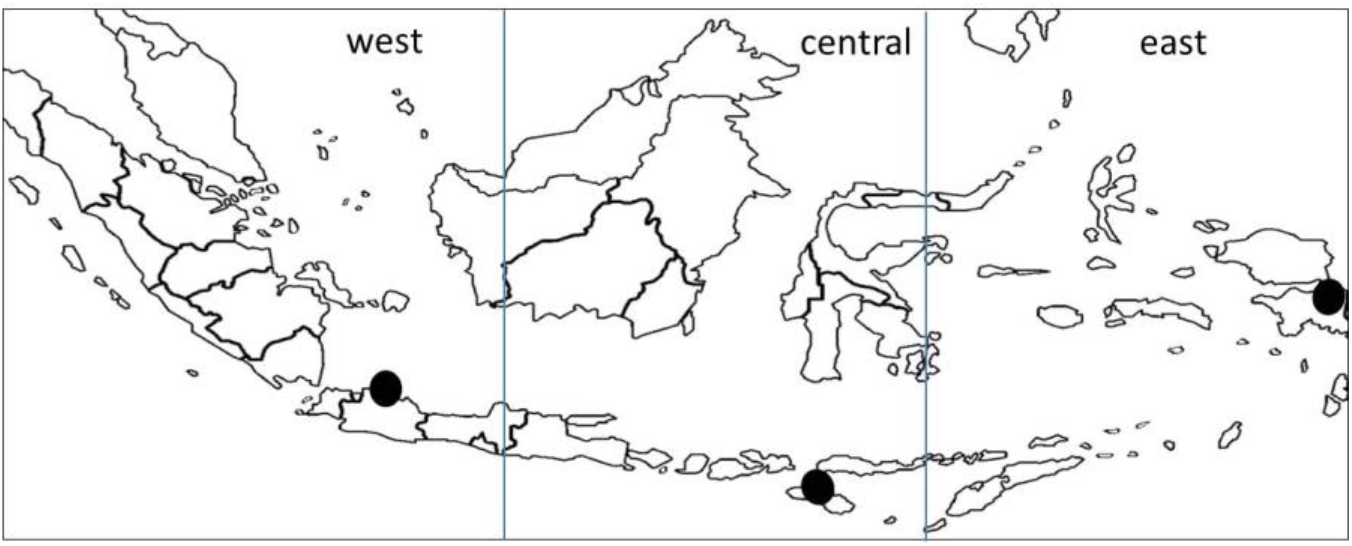
Studied locations (black dots) in west, central, and east parts of Indonesia.

### Method

#### Mosquito sampling

Mosquitos were collected as larvae or pupae and were reared in secure laboratory in artificial conditions to the adult stage. They were then frozen and stored at −80 ^°^C. The location and characteristics of each mosquito strain are available in Figure 1

#### DNA extraction

Adult specimens from each location were ground in 250 ml 10% chelex (BioRad^®^) in0.1% SDS, 1 % Tween 20,1% NP40 and the homogenate was incubated for 30 min at 56 ^°^C, 30 min at 95 ^°^C. DNA was purified by precipitation in ethanol. DNA samples were used as templates for the amplification of specific fragments of mtDNA: a 307 bp fragment for Cytb and a 597 bp fragment for cytochrome c oxidase subunit I (COI) genes (Das et al. 2016). Several sets of primers were used following (Kocher et al., 1989); for Cytb, L14841 (5’-AAAAAGCTTCCATCCAACATCTC-AGCATGATGAAA-3’) and H15149 (5’-AAACT-GCAGCCCCTCAGAATGATATTTG TCCTCA-3’). For COI, CI-J-1632 (5’-TGAT-CAAATTTATAAT-3’) and CI-N-2191 (5’-GGTA-AAATTAAAATATAAACTTC-3’).

Each reaction was performed in a Perkin-Elmer thermal cycler 2400, in a final volume of 20 μl. For Cytb and COI, the PCR mixture contained 100 ng genomic DNA, 1x buffer, 2.5 mM MgCl_2_, 250 μM each dNTP, 100 nM each primer, and 1 unit Eurobio Taq polymerase. Amplification was achieved by heating at 95 ^°^C for 5 min and then subjecting the mixture to 35 cycles of 97 ^°^C for 30 s, annealing temperature (50 ^°^C for Cytb and 40 ^°^C for COI) for 45 s, and 72 ^°^C for 1 min. The mixture was then subjected to a final extension step at 72 ^°^C for 5 min. Agarose gel electrophoresis was used to separate PCR and then purified using the Qiaquick gel extraction kit (Qiagen). Purified DNA fragments (100 ng) were directly sequenced in an automated DNA sequencer using the dideoxynucleotide-chain-termination method with ddNTPs labelled with a specific fluorochrome. After that sequences were assembled and aligned.

#### Phylogeographic analyses based on mitochondrial sequences

The mitochondrial mtDNA sequence based phylogeographic analyses for west, central, and east specimens was following methods of Tu et al. (2017) and Bayesian inference based on Huelsenbeck & Ronquist (2001), Ronquist & Huelsenbeck (2003) and Ronquist et al. (2012). In a Bayesian analysis, inferences of phylogeny are based upon the posterior probabilities of phylogenetic trees. The posterior probability of the i_th_ phylogenetic tree (τi) conditional on an alignment of DNA sequences (X) can be calculated using Bayes theorem as follows:

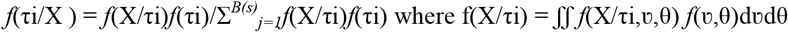

With the summation is over all *B(s)* trees that are possible for s species *[B(s)* = (2s-5)!/2^s-3^(s-3)! for unrooted trees and *B(s)* = (2s-3)!/2^s-2^(s-2)! for rooted trees], and integration is overall combinations of branch lengths (ν) and substitution parameters (θ). The prior for phylogenetic trees is *f*(τi) and is usually set to *f*(τi) = 1/ *B(s).* The prior on branch lengths and substitution parameters is denoted as *f*(ν,θ). Typically, the likelihood function *f*(X/τi,ν,θ) is calculated under the assumption that substitutions occur according to a time-homogeneous Poisson process. The same models of DNA substitution used in maximum likelihood analyses can be used in a Bayesian analysis of phylogeny. The summation and integrals required in a Bayesian analysis cannot be evaluated analytically and it uses Markov chain Monte Carlo (MCMC) to approximate the posterior probabilities of trees. MCMC is a method for taking valid, albeit dependent, samples from the probability distribution of interest, in this case the posterior probabilities of phylogenetic trees.

#### RSCU and CAI

Relative synonymous codon usage (RSCU) and Codon Adaptation Index (CAI) were calculated following methods by Liu et al. (2017), Priyono et al. (2020), and Ibis (2020). The CAI can be used to estimate gene expression levels.

Figure 2 presents the phylogeography of *A. albopictus* using samples from Asian continent as the out groups. By comparing the out groups, it is clear that *A. albopictus* populations in Indonesia have performed a distinct clade separated from the Asian continent. While in Indonesia, the *A. albopictus* populations were diversified into 2 distinct clades. First clade is grouping the west and central parts as originated from the single ancestors since they have mtDNA similarity. While there was a distinct population as can be observed in east part of Indonesia. West and central populations show that those populations were originated from the same ancestors. Whereas the sampling locations in west and central were separated by the presences of sea and island. A possible *A. albopictus* dispersal crossing islands is can be facilitated through anthropogenic assisted dispersals. According to Salgueiro et al. (2019), the airports and airlines play important role in the spread of vector-borne diseases and it has been helpful in predicting the risks of vector-borne disease importation and establishment. This can be seen in Dengue cases in Cape Verde that is in a strategic route linking Africa and South America with the major Dengue outbreak in Cape Verde was caused by a Dengue virus that originated from Senegal. This can happen since Cape Verde received more than 7,000 travelers from zika virus-affected countries, including direct flights from Brazil. In our study, the importation of *A. albopictus* from west to central parts was through anthropogenic movement considering that central parts have numerous tourism sites that frequently were visited by visitors from west parts. The similarity of mtDNA of *A. albopictus* in 2 locations was mostly due to spreading via human activities (Mousson et al. 2005, Goubert et al. 2016).

**Figure 2.**
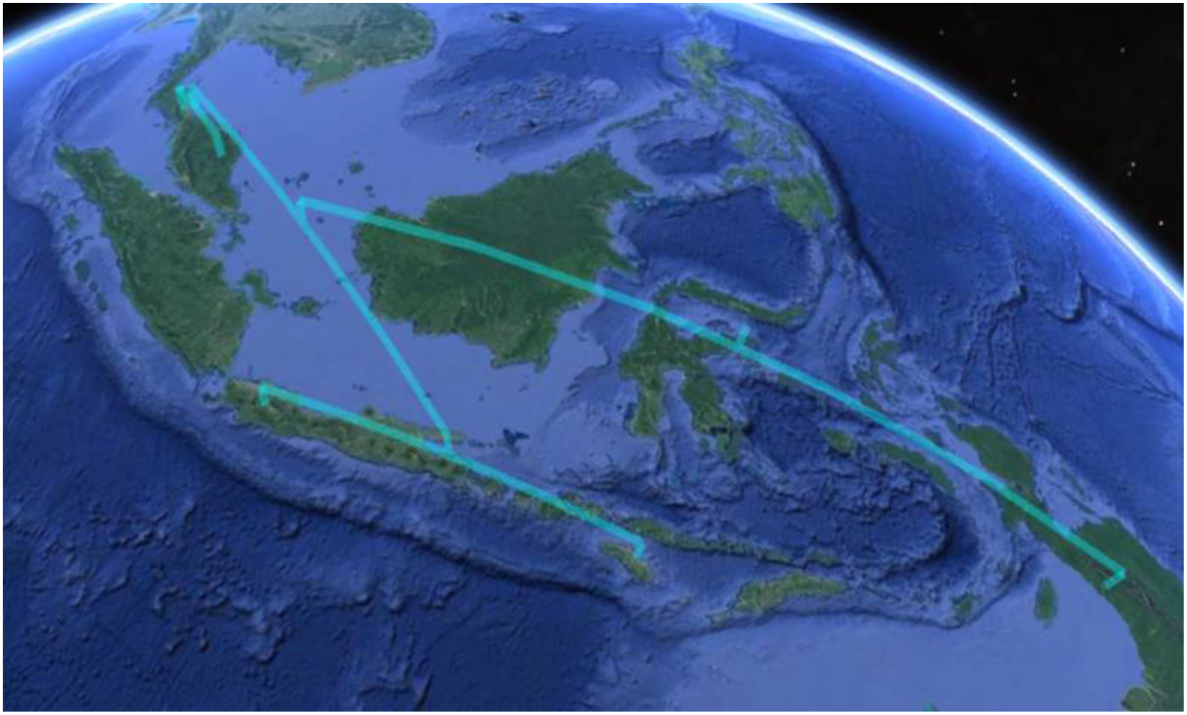
Phylogeography of *A. albopictus* DNA in Asia continent (out group), west, central, and east parts of Indonesia based on Bayesian Inference.

Relative synonymous types and usage value (RSCU) of the studied genome of *A. albopictus* is summarized in Figure 3. In *A. albopictus* from west and east parts, the codons TTG, CTG, GTG, ACG, GCG, and GGG were seldom represented. While GGA, CGA, and followed by TAT, CAT, CCT, TCT, AAC, and TTC codons were observed to be higher. The differences between west and east parts can be observed in proline amino acid signaled by CCT and CCC codons. In this amino acid, west *A. albopictus* has higher CCT than CCC. While for east mosquito, CCT and CCC were equal. To understand the pattern of non-random usage of synonymous codons in these *A. albopictus*, relative synonymous codon usage (RSCU) of individual codons were compared among *A. albopictus* populations from 3 locations. RSCU > 1 represents codons that are used more frequently than expected whereas RSCU < 1 represents codons which are used less frequently than expected. The expected number is the total number codons divided by the number of synonymous codons of each amino acid. The number of frequently used codons and rarely used codons are similar among 3 locations (Table 1). It was found a total of 60 codons with 40% codon have RSCU > 1, 21.6 % has RSCU < 1, and the rest codons were not used. In this study, Glu, Gly, Arg, and followed by Tyr, His, Pro, Ser, Asn, and Asp were the most common amino acids. This finding is corroborated with the findings of Behura and Severson (et al. 2012). Among hymenoptera that where mosquito belongs to, Arg and Asn were amino acids commonly used with high frequencies and RSCU > 1.

**Figure 3.**
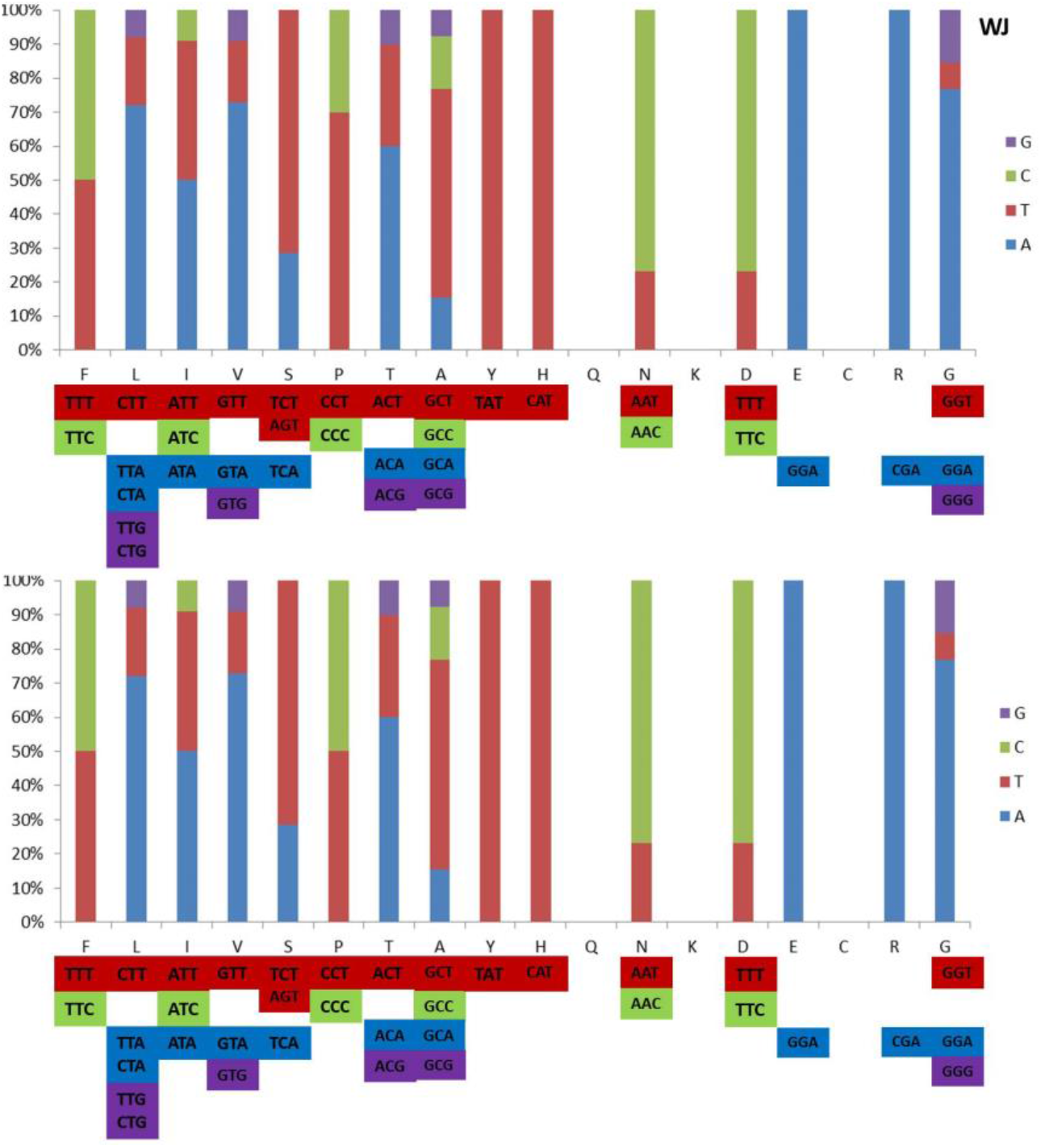
Relative synonymous codon usage (RSCU) of *A. albopictus* mtDNA in west (top) and east (bottom) parts of Indonesia (amino acid code F = Phe, L = Leu, I = Ile, V = Val, S = Ser, P = Pro, T = Thi, A = Ala, Y = Tyr, H = His, Q = Gln, N = Asn, K = Lys, D = Asp, E = Glu, C = Cys, R = Arg, G = Gly).

**Table 1.** Codons that are used more frequently than expected (RSCU >1) among the *A. albopictus* mtDNA of COI gene in west and east parts of Indonesia.

| RSCU | West |  | Central |  | East |  |
| --- | --- | --- | --- | --- | --- | --- |
|  | #codon | % | #codon | % | #codon | % |
| <1 | 13 | 21.6 | 13 | 21.6 | 13 | 21.6 |
| >1 | 24 | 40 | 24 | 40 | 24 | 40 |

In this study, CCC codon that signals proline (Pro) amino acid in east parts has higher RSCU than in west parts. In *A. albopictus*, proline plays a unique role in supporting both the flight and post-blood meal physiology of *Aedes*. Proline supports the proline-alanine shuttle system in transporting acetyl units, in the form of proline, from the fat body and to the flight muscles. In this way, proline serves as an energetic substrate for flight metabolism. The enzyme alanine aminotransferase (ALAT) is a key component of this shuttle system that enables transferring of proline and alanine as well as the detoxification of ammonia (Scaraffia & Wells 2003, Scaraffia et al. 2005, Rivera-Pérez et al. 2017).

The differences of phylogeography between *A. albopictus* in west and east parts are corresponded to proline amino acid difference and this can be related to the geographical conditions and landascape of west and east parts where the samples were collected. The landscapes in west parts were more flat in comparison to the landscape in east parts (Figure 4). The east parts were dominated by hill and landscape. This condition will affect and shape the speciation of the particular *Aedes* species. For aerial animal, rough landscape is related to the flight barrier and this will increase the energy demands for flights (Sapir et al. 2011).

**Figure 4.**
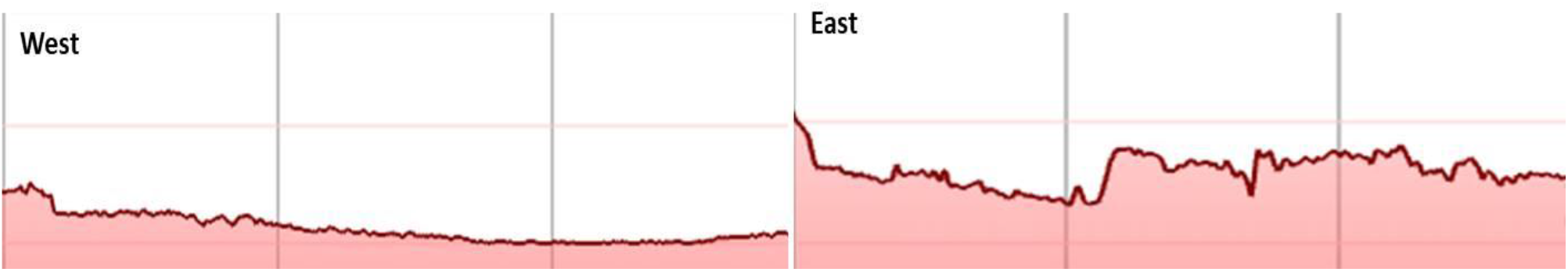
Landscapes in west and east parts of Indonesia.

Inhabiting east part with its rugged landscape is demanding flight energy and this is also applied to *A. albopictus* population in this area. Proline amino acid that distinguishes the east populations is relevant to this situation. For hymenopterans, proline is a fuel to power flight that was initially associated with the protein-rich meal of blood-sucking insects such as the tsetse fly and more recently mosquitoes. Hymenopterans use proline as a metabolic fuel to power flight is related to the unique properties of this amino acid. First, mosquito can store fuels at high concentrations in their haemolymph. Mosquito are also distinct in the high level of free circulating amino acids, where proline is predominant in many species. The high solubility of proline makes it a readily available fuel in the flight muscle and haemolymph, and does not necessitate any specific carrier protein. Proline can also serve as a carbon shuttling molecule between lipid reserves in the fat body and flight. Since inhabiting east parts requires energy to deal with the rugged landscapes, then *A. albopictus* populations in here have differences in its nucleotide sequence characterized by proline amino acid.

